# A novel paradigm for auditory discrimination training with social reinforcement in songbirds

**DOI:** 10.1101/004176

**Authors:** Kirill Tokarev, Ofer Tchernichovski

## Abstract

Zebra finches are a highly social, gregarious, species and eagerly engage in vocal communication. We have developed a training apparatus that allows training zebra finches to discriminate socially reinforced and aversive vocal stimuli. In our experiments, juvenile male zebra finches were trained to discriminate a song that was followed by a brief air puff (aversive) and a song that allowed them to stay in visual contact with another bird, ‘audience’ (social song). During training, the birds learned quickly to avoid air puffs by escaping the aversive song within 2 sec. They escaped significantly more aversive songs than socially reinforced ones, and this effect grew stronger with the number of training sessions. Therefore, we propose this training procedure as an effective method to teach zebra finches to discriminate between different auditory stimuli, which may also be used as a broader paradigm for addressing social reinforcement learning. The apparatus can be built from commercially available parts, and we are sharing the controlling software on our website.

---

Zebra finch (*Taeniopygia guttata*) is a highly social, gregarious, species that lives in colonies with high mobility of members but monogamous pairs (Adkins-Regan, 2002; Goodson et al., 2012; Zann, 1994). Zebra finches use various vocal signals, for communication: multiple call types (Marler, 2004) and a male song, which varies slightly depending on whether it is directed to another individual or not (Jarvis et al., 1998; Sossinka and Böhner, 1980). There are multiple parallels between speech acquisition in humans and vocal imitation in songbirds (Bolhuis et al., 2010) at the molecular (Fisher and Scharff, 2009), neuroanatomical (Jarvis, 2004) and behavioral levels (Doupe and Kuhl, 1999; Lipkind et al., 2013). Social aspect of vocal imitation in zebra finches attracts a lot of interest, but only few studies have addressed it directly. In accordance with the idea that individual recognition must be important in social groups of zebra finches, it is known that there is a preference towards mate’s song in females (Clayton, 1988) and father’s song in his offspring (Riebel et al., 2002), for example. Here, we propose a method that allows to study discrimination of songs and other auditory stimuli in zebra finches.

We exploited eagerness of zebra finches to engage in social communication (Goodson et al., 2012) as means for positive reinforcement in our training paradigm. We developed a social-reinforcement apparatus where birds can interact through a small window (Fig. 1).

**Figure 1.**
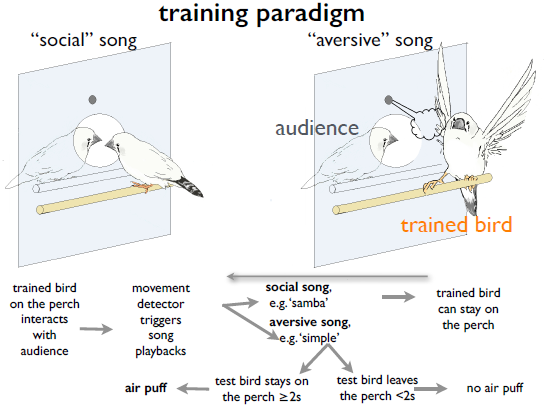
Social reinforcement apparatus: playbacks of the “social” song allow birds to socialize via a window; playbacks of the “aversive” song are followed by an air puff, which the bird learns to avoid. Artwork courtesy of Lotem Tchernichovski.

Our system is similar to existing go-nogo song discrimination boxes (Cynx and Nottebohm, 1992; Scharff et al., 1998), but instead of food reward we offer the bird a social reward: the apparatus includes two chambers and a window, through which they can communicate. When our system detects the bird perching next to the window, it sparsely plays songs. One song has no consequences, allowing the birds to keep interacting (social song). The other song is followed by an air puff (aversive song), which the bird can prevent by escaping within 2 seconds.

We use a custom-built apparatus for these experiments. Electric air valve solenoid (Assured Automotive Company) with a 12V AC power source delivers pressurized air through a hose. A sensor (SB12, Banner Engineering Corp.) uses invisible infra-red beam to detect presence of the bird at the perch next to the window between the chambers of the cage. These are connected to a PC via NI USB-6501 card (National Instruments Corp.), which in turn sends out the audio via a loudspeaker set at 80db. OT wrote a software, “Bird Puffer”, that controls this system and allows changing such parameters as length and delay of the air puff, odds for playbacks, selection of particular audio files etc. It automatically records each instance of the playback (social or aversive), and if the bird escaped it or not (within the time limit preset by the user). Bird Puffer is freely available on our website http://ofer.sci.ccny.cuny.edu

During our pilot experiments, we used 10 pairs of male juvenile zebra finches kept in isolation except for the time of the training. One animal in each pair was trained in the compartment with air puffs, and the other served as the audience. The animals spent most of their time next to the window and in most cases learnt to discriminate songs within three one-hour sessions, as they escaped significantly more aversive songs than socially reinforced ones (Fig. 2, left). The songs used for this experiment were recorded previously from two different birds, and their role (social vs aversive) was counterbalanced for different pairs. The difference between escape rates for the aversive and social songs only increased with the number of sessions, as the birds kept on escaping more and more aversive songs but not social ones (see example on Fig. 3). Escape rate from the aversive song was significantly higher in the last five sessions than in the first three, and in both cases it was higher than escape rates from the social song (p<0.005, repeated measures ANOVA, pairwise comparisons; Fig. 2).

**Figure 2.**
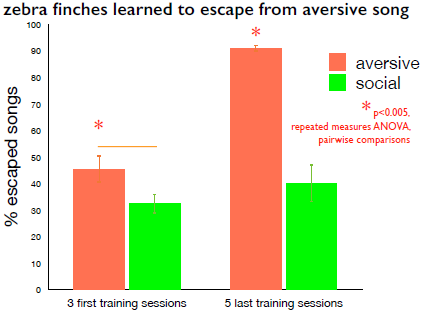
Zebra finches learned to escape from aversive song: Escape rates for the aversive song were significantly higher than for the social song even in the first three training sessions across 10 birds. In the last 5 sessions, this difference became even bigger, as the escape rate for the aversive song increased significantly compared to the first.

**Figure 3.**
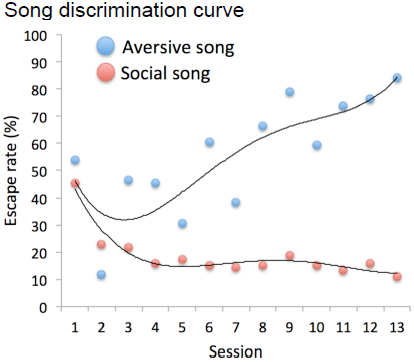
Discrimination curve for an individual bird: Escape rates for the aversive song increased with the number of training sessions but decreased for the social song.

Thus, we provide an effective social reinforcement training paradigm for zebra finches, which we expect to be an important tool in understanding the social aspect of vocal imitation.

